# Imaging chromatin interactions at sub-kilobase resolution Via Tn5-FISH

**DOI:** 10.1101/601690

**Authors:** Xu Zhang, Jing Niu, Guipeng Li, Qionghai Dai, Dayong Jin, Juntao Gao, Michael Q. Zhang

## Abstract

There is increasing interest in understanding how the three-dimensional organization of the genome is regulated. Different strategies have been employed to identify chromatin interactions genome wide. However, due to the current limitations in resolving genomic contacts, visualization and validation of these genomic loci with sub-kilobase resolution remain the bottleneck for many years. Here, we describe Tn5 transposase-based Fluorescence *in situ* Hybridization (Tn5-FISH), a Polymerase Chain Reaction (PCR)-based, cost-effective imaging method, which achieved the co-localization of genomic loci with sub-kilobase resolution, to fine dissect genome architecture at sub-kilobase resolution and to verify chromatin interactions detected by Chromatin Configuration Capture (3C)-derivative methods. Especially, Tn5-FISH is very useful to verify short-range chromatin interactions inside of contact domain and Topologically Associated Domain (TAD). It also offers one powerful molecular diagnosis tool for clinical detection of cytogenetic changes in cancers.

## Introduction

Chromatin interactions play essential roles in three-dimensional organization of the eukaryotic genome, a process critical to vital cellular functions such as transcriptional regulation^1,2^. Several strategies were employed to study genome 3D structure^3–5^: One is genome mapping techniques, including ligation-based and ligation-free methods; another is *in situ* imaging of DNA, RNA, and protein in the nucleus; and the last one is computational methods to predict either chromatin interactions or 3D chromatin organization^6^. Among these strategies, 3C-derivative methods rely on digestion and proximity ligation to capture pair-wise^5,7^ or multiple chromatin contacts^8^, while ligation-free methods, such as Genome Architecture Mapping (GAM)^9^, Split-Pool Recognition of Interactions by Tag Extension (SPRITE)^10^, and ChIA-Drop^11^ can detect multiple chromatin interactions genome-wide, by demonstrating that multiple enhancers and highly transcribed regions are associated simultaneously.

As Complementary approaches to these molecular mapping methods, fluorescence *in situ* hybridization (FISH) utilizes plasmids or PCR fragments that contain certain region of genome, to amplify and label fluorescence-tagged probes for hybridization. Multiplexed super-resolution FISH^12,13^ and different orthogonal CRISPR–dCas9 systems^14,15^ can be employed to label DNA at large scale and to identify cooperative higher-order chromatin interactions. With super-resolution^16–21^ and even electron microscopy with higher resolution^22^, more and more detailed genomic features about regulatory architecture such as chromatin loops, TADs and contact domains could be investigated.

On the other hand, chromatin interactions, either identified by 3C-based technologies or predicted by new designed algorithms, require rapid experimental verification by either 3C or imaging methods^23^. However, due to the current limitations of traditional FISH in resolving genomic contacts, validation of these genomic contacts remains the bottleneck for investigating the regulation of genome architecture. For example, many chromatin loops are inside of a TAD or contact domain^2^ (0.2-1 Mb), the basic unit of genome 3D structure and gene regulation, but typical BAC clones expand 100-300 Kb^24^, which is even comparable with the length of a TAD. Therefore, the genomic resolution of traditional FISH based on BAC clones is too low to precisely label and to image the interactions within the genomic distance less than 500 Kb, such as inside of a TAD or contact domain, whereas most of the chromatin interactions identified by 3C-based techniques fall into this range^25–27^.

There are advanced FISH techniques with higher genomic resolution, such as oligopaint^28^, HD-FISH^29^, CasFISH^30^ and MB-FISH^31^, etc. But they are either very expensive^28,31^ or require complex preparation^29,30^, which is difficult for laboratories that do not routinely use various FISH techniques to adopt and to study genomic loci. Molecular beacon-based FISH^31^ can offer a resolution of 2.5 kb, but is also technically complex and lacks cost-effectiveness.

Here we report a novel and cost-effective method named Tn5-FISH (Tn5 transposase based fluorescence *in situ* hybridization) that offers more than one order of magnitude higher resolution than traditional FISH. Tn5-FISH utilized the hyperactive Tn5 transposase for probe library construction, and PCR for probe library amplification and labeling. Foreign DNA sequences can be inserted into host genome with the help of Tn5 transposase through “cut and paste” mechanism^32^, which granted Tn5 transposase the ability to induce the DNA double breaks at insertion site. Efficient segmentation of targeted DNA sequence is favored in probe library construction^33,34^. With the aid of bioinformatic tools, we can obtain the repeat-free DNA sequence suitable for probing via genomic PCR.

Then Tn5-FISH was employed to verify the chromatin interactions, for example, the ones inside of type II KRT locus in chromosome 12 of K562 cells measured by Hi-C contact map from published data. Triple-color Tn5-FISH can verify these interactions well, with triple-color traditional BAC FISH as positive control, indicating that Tn5-FISH is suitable to visualize specific genomic loci and chromatin interactions inside a typical TAD, in either normal cell types, cancer cell line, or even tissues. Furthermore, Tn5-FISH potentially becomes one of the best clinical tools for the rapid molecular diagnosis, to detect or to confirm classic cytogenetic changes in cancers.

## Results

### The Scheme of Tn5-FISH to label target genomic loci

Foreign DNA sequences can be inserted into host genome with the help of Tn5 transposase through “cut and paste” mechanism^32^, offering Tn5 transposase the ability to induce DNA double breaks at insertion site. Therefore, Tn5-FISH utilizes the hyperactive Tn5 Transposase for probe library construction, and PCR for probe library amplification and labeling (see the schematic diagram in **Fig. 1**. For more details, see **Supplementary Note 1**). Efficient segmentation of targeted DNA sequence is favored in probe library construction^33,34^. With the aid of UCSC Genome Browser, one could easily obtain the repeat-free DNA target sequence suitable for probing via genomic PCR.

**Figure 1.**
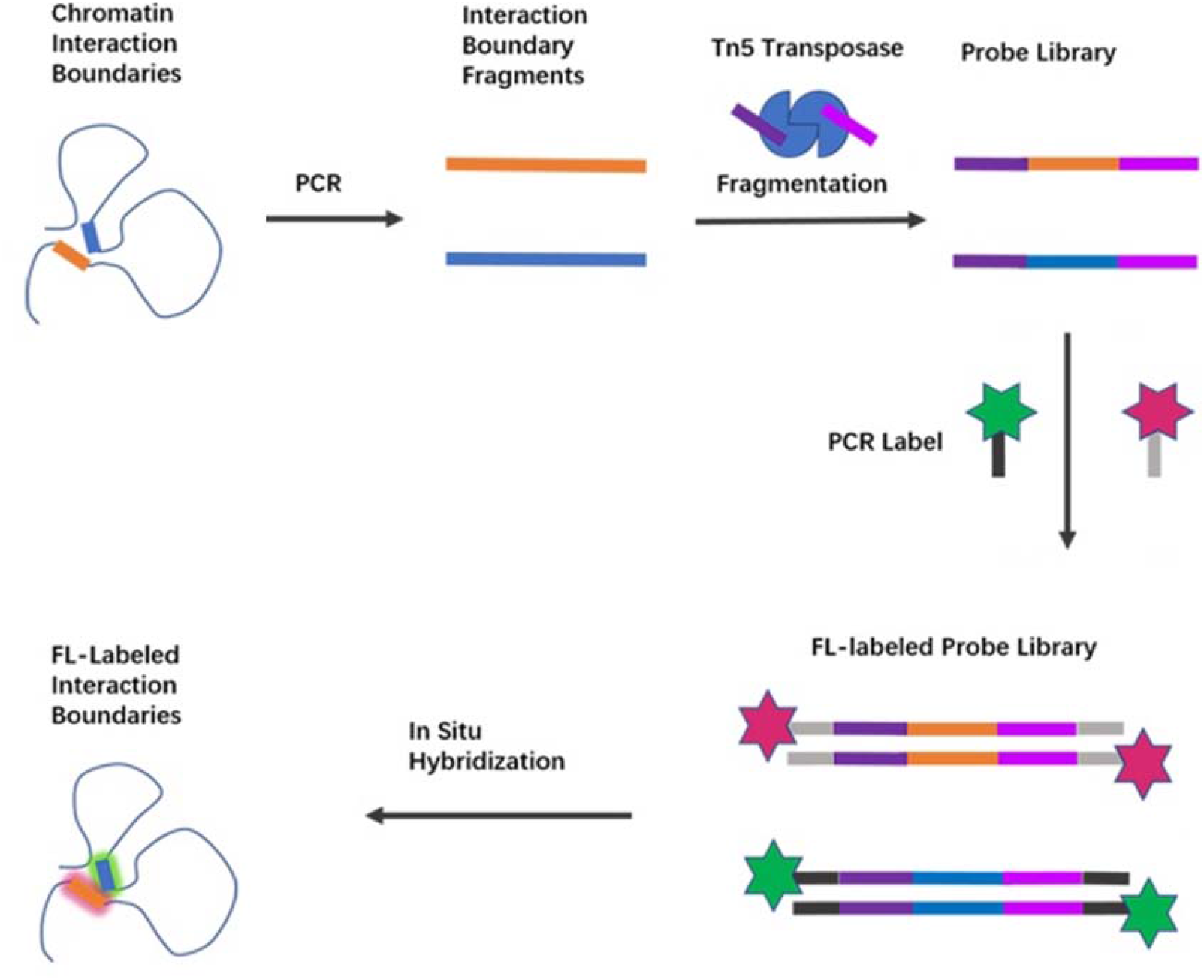
The Scheme of Tn5-FISH to label target genomic loci. The genomic loci (orange and blue) were amplified by PCR, fragmented by Tn5 transposase, from which universal tag sequence (purple and magenta) were added. Then the 2^nd^ round of PCR was used to amplify the probe and fluorescence labeling. After *in situ* hybridization, the chromatin interaction boundaries (labeled as red and green, respectively) can be labelled and imaged. FL-labeled: Fluorescence-labeled.

We next tested the specificity of Tn5-FISH by using two probes targeting two adjacent loci GM19705 and Platr22 (only 6.5 kb away from each other, **Supplementary Fig. S1a**), in wild-type mouse Embryonic Stem Cells (WT mESCs) and Platr22 knock-out mESCs (Platr22-KO mESCs, **Supplementary Fig. S1b**), respectively. As proposed, Platr22 loci were only visible in WT mESCs, but not in Platr22-KO mESCs, while GM19705 loci can be visualized in either WT or Platr22-KO mESCs (**Fig. 2a-d**). These results were simultaneously verified by traditional FISH with BAC clone (BMQ-36B16) which covered both GM19705 and Platr22 loci, indicating that Tn5-FISH is a versatile technique for labeling unique genomic loci with high specificity. A full width at half maximum (FWHM) of each fluorescent signal density, which approximately follow a Gaussian distribution, is 230 ± 20 nm and 300 ± 40 nm (mean ± SD) for the TAMRA and Alexa Fluor 488, respectively (**Fig. 2e**), suggesting a significant improvement of spatial resolution from ~300 nm for BAC FISH to ~230 nm for Tn5-FISH (**Fig. 2f**), benefited from the smaller size of probes (~4Kb).

**Figure 2.**
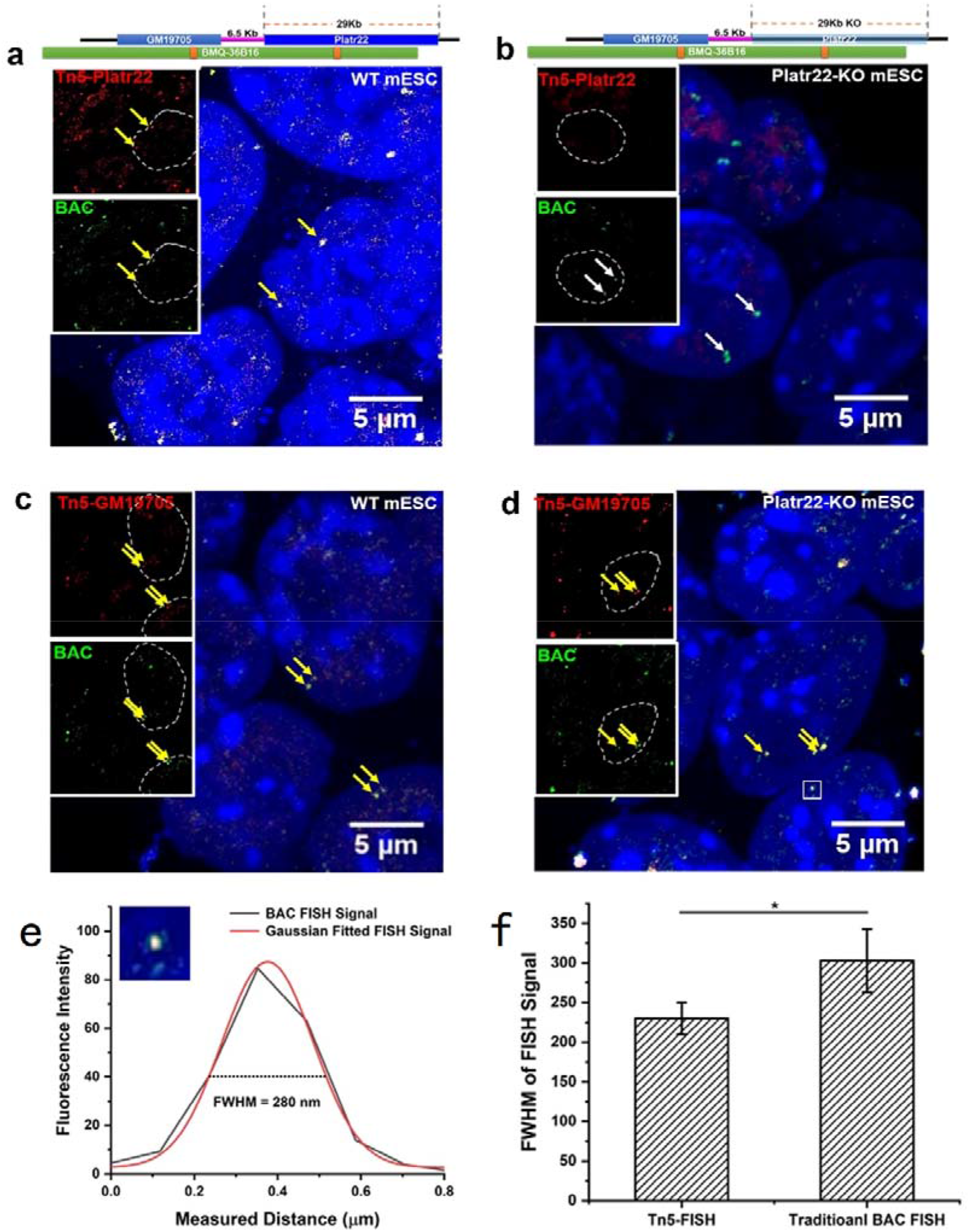
The specificity of Tn5-FISH. **a-d** The specificity of Tn5-FISH signals (red dots) was verified by traditional BAC FISH (green dots) in mESCs. In every panel, red channel and green channel are merged with DAPI staining (blue). BAC probe (green) and Tn5-Platr22 probe (red, **a** and **b**) or Tn5-GM19705 probe (red, **c** and **d**) were hybridized in mESCs (the left two panels **a** and **c**), or in Platr22-KO mESC cells (the right two panels **b** and **d**) simultaneously. Yellow arrowhead indicates co-localized FISH signals, while white arrowhead indicates no co-localization observed. All images were taken by laser confocal scanning microscopy (LCSM). **e** The distribution of fluorescence intensity from a representative FISH signal (white box in the bottom right panel in **d**), was fit to a Gaussian profile to determine its FWHM. **f** Statistical analysis (t-test) of FWHM values measured from 50 Tn5-FISH signals co-localized with BAC FISH signals, indicates that the spatial resolution of Tn5-FISH is significantly (p<0.05 as indicated by asterisk) higher than that of BAC FISH.

### Resolution of Tn5-FISH

Next we tried to test the genomic resolution limit of Tn5-FISH. With traditional FISH as reference, we gradually shortened the target genome locus fragment sizes from ~8kb downwards (7.99kb, 4.14 kb, 2.58Kb, 1.17Kb, 0.8Kb, 0.5Kb, and 0.3Kb, respectively, as shown in **Fig.3 a-g**). These seven different fragments were imaged in K562 cells with Structured Illumination Microscopy (SIM) (**Fig.3 a-g**).

**Figure 3.**
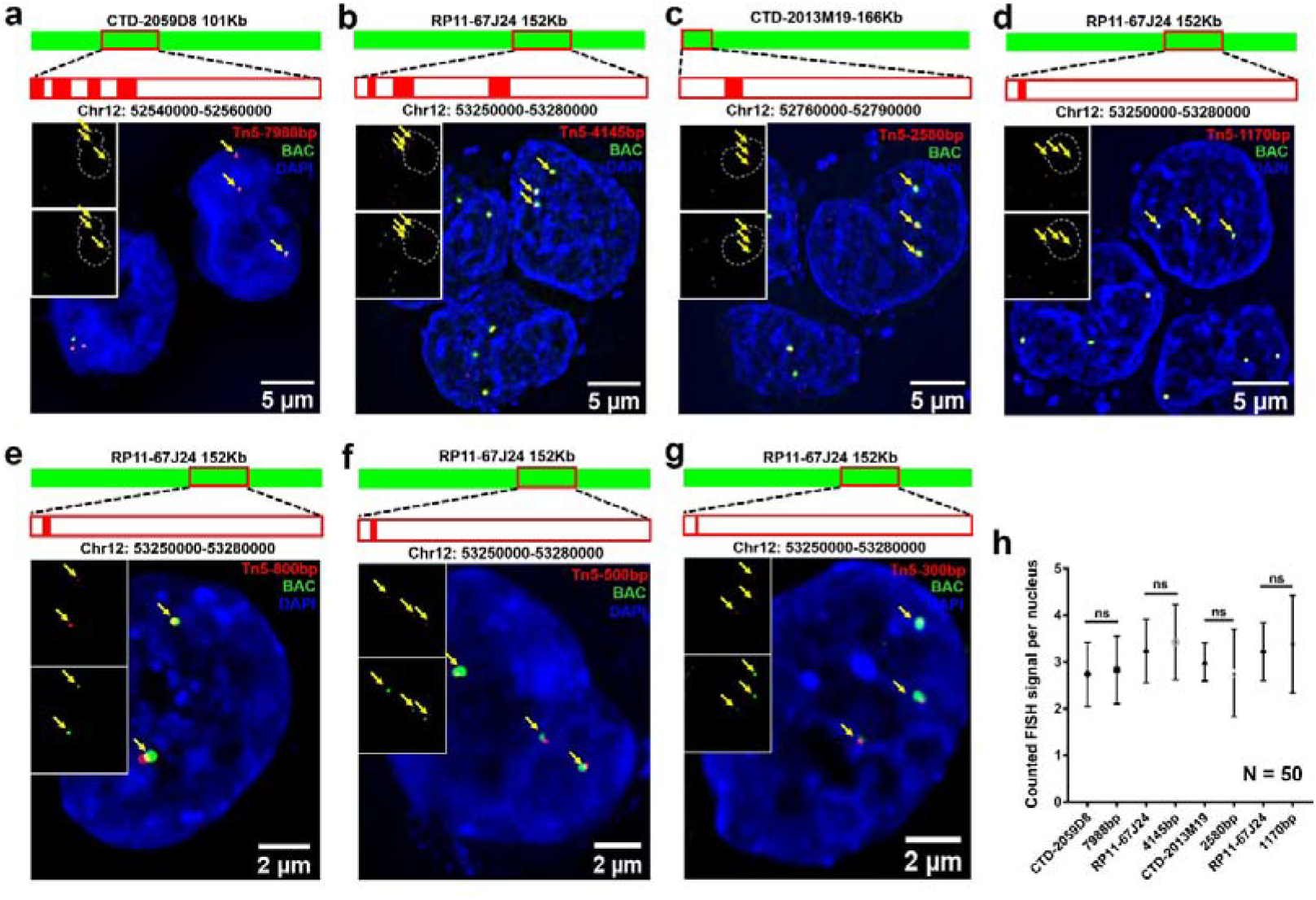
The resolution of Tn5-FISH by SIM. **a** Tn5 probes (four red bars) with genomic resolution of 7988 bp (i.e. the sum of the length of four red bars) and BAC probe with genomic resolution of 101 kb (green bar) were hybridized and co-localized in K562 cells simultaneously. Yellow arrow indicates the imaging of single locus labeled by either Tn5-FISH (red channel, top left insert) or BAC-FISH (green channel, bottom left insert). Both Tn5-FISH (labeled with TAMRA) and BAC-FISH signals (**a-d** labeled with Alexafluor488, **e-g** labeled with Alexafluor647) were merged with DAPI staining (blue). All images here are the Maximum Intensity Projection, with scale bar of 5μm. **b** same as **a**, except with Tn5 probe with genomic resolution of 4145 bp and BAC probe with 152 Kb. **c** same as **a**, except with Tn5 probe with genomic resolution of 2580 bp and BAC probe with 166 Kb. **d** same as **a**, except with Tn5 probe with genomic resolution of 1170 bp and BAC probe with 152 Kb. **e** same as **a**, except with Tn5 probe with genomic resolution of 800 bp and BAC probe with 152 Kb. **f** same as **a**, except with Tn5 probe with genomic resolution of 500 bp and BAC probe with 152 Kb. **g** same as **a**, except with Tn5 probe with genomic resolution of 300 bp and BAC probe with 152 Kb. **e**, The target genome locus fragment sizes were gradually shortened from 7988bp downwards, to 4145, 2580 and 1170bp, respectively. There is no statistical difference (t-test) observed between the hybridization efficiency of Tn5-FISH and that of BAC FISH. In each group, FISH signals in 50 Cells were counted.

Tn5-FISH signals were then quantified and compared with traditional BAC FISH respectively (**Fig. 3h**). Even with a target as short as around one kilobase, good fluorescent signal can still be observed (with 86% hybridization efficiency compared with traditional BAC FISH, **Supplementary Table S1**). Therefore, Tn5-FISH can offer a genomic resolution of sub kilobase, which is more than one order of magnitude higher than traditional FISH methods.

Furthermore, we compared the cost-effectiveness of some different FISH methods published so far^28–31,35^. With the cost of only ~50$ and only 1 day per replicate in one experiment (**Table 1**), Tn5-FISH is the most cost-effective one among various FISH methods, according to our knowledge.

**Table 1:** Comparison of Tn5-FISH with other FISH methods.

|  | **Time for probe preparation** | **The amount of DNA per batch** | **Genomic resolution** | **cost per experiment (US Dollar)** |
| --- | --- | --- | --- | --- |
| **Tn5-FISH** | ~1 day | 50 ng | 1.17 kb | 46 |
| **HD-FISH**<sup>27</sup> | ~1 week | library | 10 kb | 288 |
| **MB-FISH**<sup>29</sup> | ~1 month | 10 μM | 2.5 kb | 12,104 |
| **casFISH**<sup>28</sup> | ~1 week | 5-25 nM | 5 kb | 410 |
| **BAC FISH**<sup>33</sup> | ~2 days | 1-2 μg | 100-200 kb | 227 |
| **Oligopaints**<sup>26</sup> | ~1 week | library | 5 kb | 757 |

### Imaging the chromatin interactions with Tn5-FISH in a TAD of cancer cells

Sub-kilobase-resolution Tn5-FISH offers great potential to be one powerful tool in clinical molecular diagnosis. Therefore, we asked whether Tn5-FISH can be used for the fine dissect genome architecture in cancer cells. Different combination of keratin proteins encoded by KRT locus (keratin-encoding gene locus in one TAD), can be assembled into keratin intermediate filaments as structure scaffolds^36^. Mutations in KRT genes may cause human disease such as epidermolysis bullosa simplex and pachyonychia congenita. Chromatin interactions inside of type II KRT locus in chromosome 12 of K562 cells (**Fig. 4a**) were measured by Hi-C contact map (**Fig. 4b**) from the published data^37^ which indicates that there are multiple chromatin loops within the TAD. The chromatin loops were verified by triple-color Tn5-FISH (**Fig. 4c** and **Supplementary Figure S2**), with triple-color traditional BAC FISH as a positive control (**Supplementary Figure S3**).

**Figure 4.**
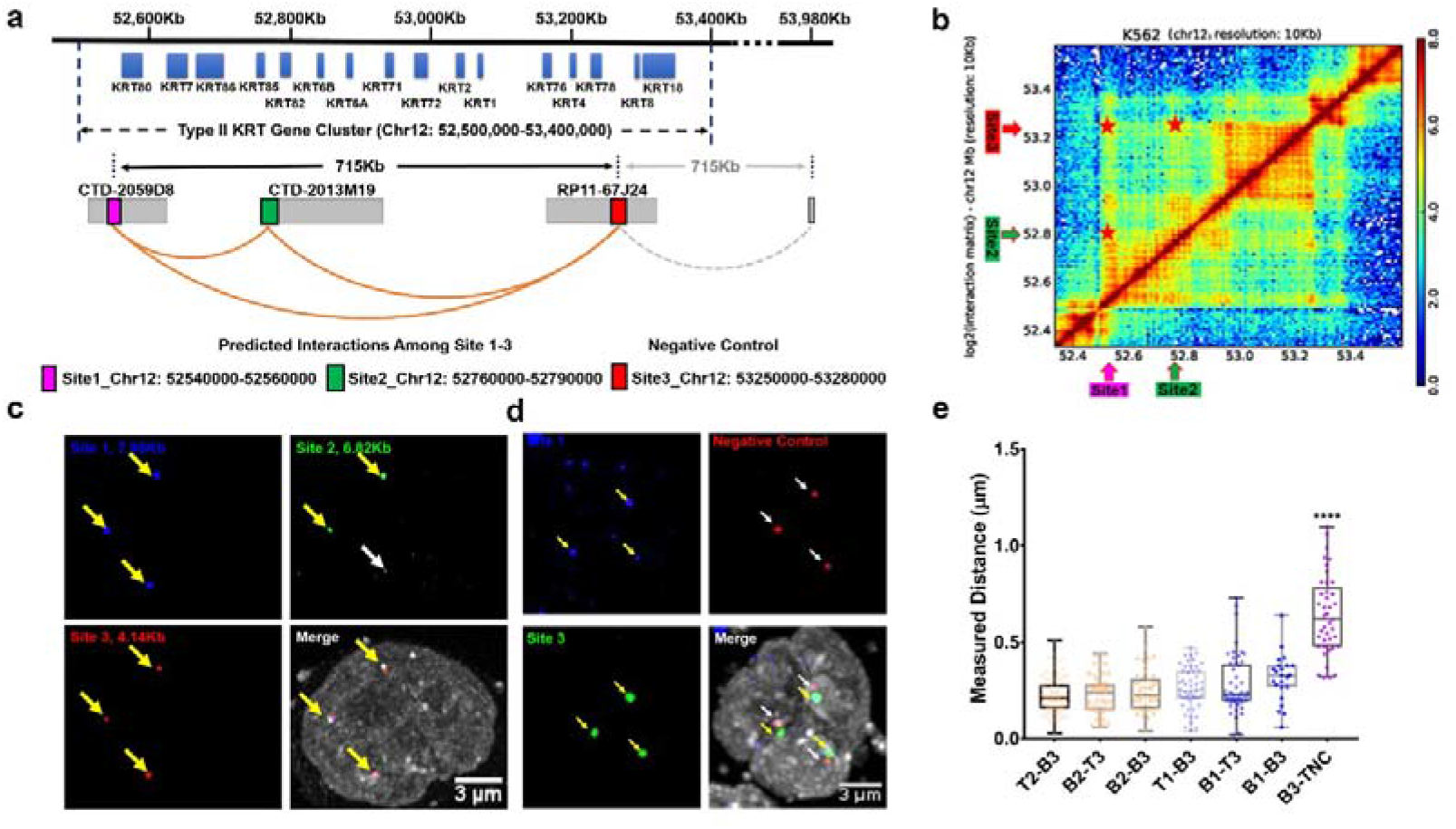
Multi-color Tn5-FISH verifies predicted chromatin interactions in K562 cells. **a** A diagram of predicted chromatin interactions (denoted by orange curves) within KRT gene cluster II in K562 cells, interpreted from (**b**) the interaction frequency matrix at Chr 12: 52500Kb-53500Kb. Site 1, 2, 3 was marked in magenta, green and red, respectively. **c** The predicted interactions among these 3 sites were verified in K562 cells. The yellow arrow indicates co-localization other two sites, and the white arrow indicates no co-localization with other two sites. The length of the probes for site 1, 2, and 3 are 7.9kb, 6.8kb and 4.1Kb, respectively. **d** The negative control indicated non-interaction spatial distance was more than 0.5 μm away from the interacting genomic loci (site 1 to site 3) with the same genomic distances. **e** The spatial distances of interacting genomic loci is statistically different from that of non-interacting loci, but for interacting loci, no statistical differences were identified between Tn5-FISH labeled or BAC FISH labeled genomic loci (one-way ANOVA).

The chromatin loops were verified by triple-color Tn5-FISH (**Fig. 4c**), with triple-color traditional BAC FISH as a positive control (data not shown). Three interacting sites were closely adjacent to each other spatially (no matter labeled by BAC FISH or by Tn5-FISH), while the negative control was ~500 nm away from the interacting sites (**Fig. 4d and e**). The independent verification by both different FISH methods further indicates that Tn5-FISH is a robust method to investigate chromatin interactions inside of a given TAD in 3D genome structure.

## Discussion

As very different strategies, both genome mapping and imaging provide very different, yet complementary in many ways, information about the chromatin looping and genome architecture. Many chromatin loops fall into the range of one BAC, thus can’t be confirmed by traditional BAC FISH. Tn5-FISH, a cost- and time-effective imaging method, expanded access to validate or reconcile the contact frequency of genomic loci in Hi-C contact map, in either normal cell types or cancer cell lines, thus is a good choice for the imaging and verification.

### Improving the specificity and reducing the uncertainty

Tn5 Transposase could generate short segments around 40nt, reaching a theoretical maximum labeling density of 50 probes per kb, therefore helps the signal to be distinguished from background. There are three reasons to use Tn5 for probes preparation in mammalian FISH. First, the hyperactive Tn5 provides the flexibility of DNA probe template (BAC or PCR fragments), in small quantities (1-50ng, can be varied according to commercially available Tn5 library construction kits), which is 20 times lesser than the amount BAC clone required. Second, the full length of probes in constructed library is normally within 100-200 bp, which is appropriate for FISH. Third, the DNA sequence from fully functional Tn5-DNA complex is orthogonal to mammalian genome^38^, therefore the specificity of probes was guaranteed.

The uncertainties of FISH experiment might arise from probe size, chromatin displacement during denaturation and hybridization, background noise and ambiguity caused by homologous chromosomes, etc^26^. Tn5-FISH method, which uses the fragments of several kilobases to generate probes for labeling, will greatly reduce the uncertainty caused by probe size. However, the uncertainty caused by the hybridization process still exist due to the denatured hybridization conditions of Tn5-FISH similar to that of traditional FISH. On the other hand, for one specific site, when probe size decreases, the mis-target rate increases (**Table 1**). In this case, the quality of Tn5-FISH images can be complemented with BAC FISH as positive control or backup.

### What could be the highest genomic resolution for Tn5-FISH theoretically, and how to image it?

With the following assumptions: (1) Nucleotide sequences of a human genome are randomly distributed; (2) The designed oligonucleotide was hybridized very efficiently to the target sequence without or with little hybridization to other similar sequences presented in the genome; (3) The single fluorescent molecule labelled on this designed oligonucleotide can be imaged very efficiently and successfully; we may conclude that theoretically, the highest genomic resolution of FISH can reach 17 nucleotides, as an ideal 17-base-long oligonucleotide should find its specific and the only position on the genome, and 4^17^ nucleotides can already cover the whole 3-billion-base-pair human genome.

However, the super-resolution imaging of the single-fluorophore labelled on the 17-base oligonucleotide can be very challenging, if not impossible. The two-dimensional localization precision of an individual fluorescent molecule is defined by the sum of photon noise, pixelation noise and background noise, as shown by^39,40^

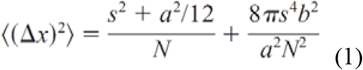

Here *S* is the standard deviation of the PSF of a given microscope system, *N* is the number of photons collected, *a* is pixel size, and *b* is background noise.

In order to obtain high-quality imaging, one needs to optimize the microscopic system to increased *s*, employ the best-performing organic fluorophore for high photon number *N*, and adopt appropriate strategy to deliver fluorophores into cell nuclei efficiently, to reduce the background noise *b*. Furthermore, one needs to consider how to tradeoff these optimized parameters. For example, during sample preparation, what labeling density and washing conditions should be employed; during imaging, what wavelength and detector parameters should be considered; whether TIRF illumination should be considered to reduce the background noise, etc. Photobleaching curve^41^ could be employed, to decide whether to increase the photon number N, or to reduce the background noise b in formular (1). According to our experience, for the single-color FISH with one labelled typical fluorophore molecule, the localization precision of less than 10nm can be obtained, with the photon number N larger than 10^4^.

In summary, Tn5-FISH can be a powerful tool for broad image-based chromatin structure studies with joint consideration of contact frequency and high-throughput bar-coding strategy. Furthermore, with much higher resolution, Tn5-FISH will potentially become one of the best clinical methods for the rapid (prenatal) diagnosis and prognosis^42,43^, to detect or confirm classic cytogenetic changes, such as chromosomal rearrangement, aberrations/abnormalities, tiny chromosomal changes in patients with Edward syndrome, Williams syndrome, hematologic malignancies, or certain solid tumors.

## MATERIALS AND METHODS

### Cell culture

K562 and GM12878 cells were purchased from China Infrastructure of Cell Line Resources (Peking Union Medical College, Beijing, China) and maintained in RPMI1640 medium (Gibco, U.S.A.) supplemented with 10% fetal bovine serum (FBS, Gibco, U.S.A.), 50 units/ml penicillin and streptomycin (Gibco, U.S.A.), and Non-Essential Amino Acids (NEAA, Gibco, U.S.A.) as instructed.

### Tn5 FISH probe preparation

Tn5-FISH probes were constructed as follows. First, sequences of interacting sites were downloaded from UCSC genome browser with repeats masked as N. DNA fragments used for probe library generation were amplified by PCR and recovered by DNA Cleanup kit (D4014, Zymo research, U.S.A.). The amplified probe library could serve as templates for Tn5-FISH probes. The Tn5-FISH probes were obtained by a second PCR amplification with fluorescence-tagged primers.

### Traditional FISH probes generated by Nick Translation of BAC clone

Traditional FISH probes were prepared as previously described^35^. Briefly, BAC clones were ordered from Thermo Fisher, and obtained by BACMAX kit (Epicenter, U.S.A.). For 1 μg BAC clone, 1 μL DNA Pol I and 20 U DNase I (NEB, U.K.), 1 mM each of dATP, dCTP, dGTP and fluorescence-tagged dUTP/dTTP were added together and incubated at 15°C for 2 h, and finally re-dissolved in DNA FISH buffer.

### Multi-color Tn5-FISH preparation

The procedure of Tn5-FISH was the same as traditional FISH as previously described^35^. Briefly, cells were fixed by 4% paraformaldehyde and permeated by 0.1% Triton-X 100, then incubated at 20% glycerol for 30 min. As for two-color Tn5-FISH, each color of 10 ng Tn5-FISH probe was mixed with DNA FISH buffer and applied to cells. The FISH program was set to 75°C for 5 min, then 37°C overnight. The slides were then imaged either by Carl Zuess LSM780 confocal microscope or Nikon A1 SIM microscope.

### Image processing and Quantification of FISH signals

The obtained Tn5-FISH images were processed by FIJI software (version 1.25h, from NIH) and the videos were processed and all images were quantified by Imaris 9.2.0 (Bitplane, Switzerland).

### Statistical analysis

The statistical analyses are carried out using build-in two-tailed t-test models provided by Graphpad Prism7 software (Graphpad Software, San Diego, U.S.A.). The significance indicators are labeled according to p-values analyzed from the data in this manuscript.

## Acknowledgement

Due to space limitations, we apologize that only a very selected list of FISH methods could be cited here. We thank Prof. Minping Qian and members from Prof. Michael Q Zhang’s lab for useful discussions. This work was supported in part by the National Natural Science Foundation of China (31671383 and 81890990), the State Key Research Development Program of China (2017YFA0505503 and 2016YFC1200300). This work was supported by funds from Beijing Advanced Innovation Center for Structural Biology, Tsinghua University.

## Reference

1 Dekker, J. & Mirny, L. The 3D Genome as Moderator of Chromosomal Communication. Cell 164, 1110–1121, doi: 10.1016/j.cell.2016.02.007 (2016).

2 Yu, M. & Ren, B. The Three-Dimensional Organization of Mammalian Genomes. Annu Rev Cell Dev Biol 33, 265–289, doi: 10.1146/annurev-cellbio-100616-060531 (2017).

3 Ma, T. et al. Developing novel methods to image and visualize 3D genomes. Cell biology and toxicology 34, 367–380, doi: 10.1007/s10565-018-9427-z (2018).

4 Gall, J. G. The origin of in situ hybridization – A personal history. Methods 98, 4–9, doi: 10.1016/j.ymeth.2015.11.026 (2016).

5 Sigal, Y. M., Zhou, R. & Zhuang, X. Visualizing and discovering cellular structures with super-resolution microscopy. Science 361, 880–887, doi: 10.1126/science.aau1044 (2018).

6 Rowley, M. J. et al. Evolutionarily Conserved Principles Predict 3D Chromatin Organization. Mol Cell 67, 837–+, doi: 10.1016/j.molcel.2017.07.022 (2017).

7 Gall, J. G. The origin of in situ hybridization – A personal history. Methods 98, 4–9, doi: http://dx.doi.org/10.1016/j.ymeth.2015.11.026 (2016).

8 Allahyar, A. et al. Enhancer hubs and loop collisions identified from single-allele topologies. Nature Genetics 50, 1151–+, doi: 10.1038/s41588-018-0161-5 (2018).

9 Beagrie, R. A. et al. Complex multi-enhancer contacts captured by genome architecture mapping. Nature 543, 519–+, doi: 10.1038/nature21411 (2017).

10 Quinodoz, S. A. et al. Higher-Order Inter-chromosomal Hubs Shape 3D Genome Organization in the Nucleus. Cell 174, 744–757.e724, doi: 10.1016/j.cell.2018.05.024 (2018).

11 Zheng, M. et al. Multiplex chromatin interactions with single-molecule precision. Nature, doi: 10.1038/s41586-019-0949-1 (2019).

12 Allahyar, A. et al. Enhancer hubs and loop collisions identified from single-allele topologies. Nature genetics 50, 1151–1160, doi: 10.1038/s41588-018-0161-5 (2018).

13 Bintu, B. et al. Super-resolution chromatin tracing reveals domains and cooperative interactions in single cells. Science (New York, N. Y.) 362, doi: 10.1126/science.aau1783 (2018).

14 Chen, B. H., Guan, J. & Huang, B. Imaging Specific Genomic DNA in Living Cells. Annu Rev Biophys 45, 1–23, doi: 10.1146/annurev-biophys-062215-010830 (2016).

15 Ma, H. H. et al. Multiplexed labeling of genomic loci with dCas9 and engineered sgRNAs using CRISPRainbow. Nat Biotechnol 34, 528–530, doi: 10.1038/nbt.3526 (2016).

16 Beliveau, B. J. et al. Single-molecule super-resolution imaging of chromosomes and in situ haplotype visualization using Oligopaint FISH probes. Nature Communications 6, doi: ARTN714710.1038/ncomms8147 (2015).

17 Bintu, B. et al. Super-resolution chromatin tracing reveals domains and cooperative interactions in single cells. Science 362, 419–+, doi: ARTNeaau178310.1126/science.aau1783 (2018).

18 Cardozo Gizzi, A. M. et al. Microscopy-based chromosome conformation capture enables simultaneous visualization of genome organization and transcription in intact organisms. bioRxiv, 434266, doi: 10.1101/434266 (2018).

19 Kundu, S. et al. Polycomb Repressive Complex 1 Generates Discrete Compacted Domains that Change during Differentiation. Mol Cell 65, 432–+, doi: 10.1016/j.molcel.2017.01.009 (2017).

20 Szabo, Q. et al. TADs are 3D structural units of higher-order chromosome organization in Drosophila. Sci Adv 4, doi: ARTNeaar808210.1126/sciadv.aar8082 (2018).

21 Nir, G. et al. Walking along chromosomes with super-resolution imaging, contact maps, and integrative modeling. PLOS Genetics 14, e1007872, doi: 10.1371/journal.pgen.1007872 (2018).

22 Ou, H. D. et al. ChromEMT: Visualizing 3D chromatin structure and compaction in interphase and mitotic cells. Science (New York, N. Y.) 357, doi: 10.1126/science.aag0025 (2017).

23 Fudenberg, G. & Imakaev, M. FISH-ing for captured contacts: towards reconciling FISH and 3C. Nature methods 14, 673–678, doi: 10.1038/nmeth.4329 (2017).

24 Shizuya, H. & Kouros-Mehr, H. The development and applications of the bacterial artificial chromosome cloning system. The Keio Journal of Medicine 50, 26–30, doi: 10.2302/kjm.50.26 (2001).

25 de Wit, E. & de Laat, W. A decade of 3C technologies: insights into nuclear organization. Genes & development 26, 11–24, doi: 10.1101/gad.179804.111 (2012).

26 Giorgetti, L. & Heard, E. Closing the loop: 3C versus DNA FISH. Genome biology 17, 215, doi: 10.1186/s13059-016-1081-2 (2016).

27 Williamson, I. et al. Spatial genome organization: contrasting views from chromosome conformation capture and fluorescence in situ hybridization. Genes & development 28, 2778–2791, doi: 10.1101/gad.251694.114 (2014).

28 Beliveau, B. J. et al. Versatile design and synthesis platform for visualizing genomes with Oligopaint FISH probes. Proc Natl Acad Sci USA 109, 21301–21306, doi: 10.1073/pnas.1213818110 (2012).

29 Bienko, M. et al. A versatile genome-scale PCR-based pipeline for high-definition DNA FISH. Nat Methods 10, 122–124, doi: 10.1038/nmeth.2306 (2013).

30 Deng, W. L., Shi, X. H., Tjian, R., Lionnet, T. & Singer, R. H. CASFISH: CRISPR/Cas9-mediated in situ labeling of genomic loci in fixed cells. Proceedings of the National Academy of Sciences of the United States of America 112, 11870–11875, doi: 10.1073/pnas.1515692112 (2015).

31 Ni, Y. X. et al. Super-resolution imaging of a 2.5 kb non-repetitive DNA in situ in the nuclear genome using molecular beacon probes. Elife 6, doi: ARTNe2166010.7554/eLife.21660 (2017).

32 Reznikoff, W. S. in Annual review of genetics Vol. 42 Annual Review of Genetics 269–286 (Annual Reviews, 2008).

33 Rich, J. J. & Willis, D. K. A SINGLE OLIGONUCLEOTIDE CAN BE USED TO RAPIDLY ISOLATE DNA-SEQUENCES FLANKING A TRANSPOSON TN5 INSERTION BY THE POLYMERASE CHAIN-REACTION. Nucleic acids research 18, 6673–6676, doi: 10.1093/nar/18.22.6673 (1990).

34 Strasbaugh, L. D., Bourke, M. T., Sommer, M. T., Coon, M. E. & Berg, C. M. PROBE MAPPING TO FACILITATE TRANSPOSON-BASED DNA SEQUENCING. Proceedings of the National Academy of Sciences of the United States of America 87, 6213–6217, doi: 10.1073/pnas.87.16.6213 (1990).

35 Bayani, J. & Squire, J. A. Fluorescence In Situ Hybridization (FISH). Current Protocols in Cell Biology 23, 22.24.21–22.24.52, doi: 10.1002/0471143030.cb2204s23 (2004).

36 Homberg, M. & Magin, T. M. Beyond expectations: novel insights into epidermal keratin function and regulation. Int Rev Cell Mol Biol 311, 265–306, doi: 10.1016/B978-0-12-800179-0.00007-6 (2014).

37 Rao, S. S. P. et al. A 3D Map of the Human Genome at Kilobase Resolution Reveals Principles of Chromatin Looping. Cell 159, 1665–1680, doi: 10.1016/j.cell.2014.11.021 (2014).

38 Gorbacheva, T., Quispe-Tintaya, W., Popov, V. N., Vijg, J. & Maslov, A. Y. Improved transposon-based library preparation for the Ion Torrent platform. Biotechniques 58, 200–202, doi: 10.2144/000114277 (2015).

39 Bobroff, N. Position measurement with a resolution and noise-limited instrument. 57, 1152–1157, doi: 10.1063/1.1138619 (1986).

40 Thompson, R. E., Larson, D. R. & Webb, W. W. Precise Nanometer Localization Analysis for Individual Fluorescent Probes. Biophysical Journal 82, 2775–2783, doi: https://doi.org/10.1016/S0006-3495(02)75618-X (2002).

41 Gordon, M. P., Ha, T. & Selvin, P. R. Single-molecule high-resolution imaging with photobleaching. Proc. Natl Acad. Sci. U. S. A. 101, 6462, doi: 10.1073/pnas.0401638101 (2004).

42 Guo, Y. et al. Multiscale Modeling of Inflammation-Induced Tumorigenesis Reveals Competing Oncogenic and Oncoprotective Roles for Inflammation. Cancer Res 77, 6429–6441, doi: 10.1158/0008-5472.Can-17-1662 (2017).

43 Guo, Y. et al. Network-Based Combinatorial CRISPR-Cas9 Screens Identify Synergistic Modules in Human Cells. ACS synthetic biology 8, 482–490, doi: 10.1021/acssynbio.8b00237 (2019).

