## Supplementary Figures and Information for "Imaging chromatin interactions at sub-kilobase resolution Via Tn5-FISH"

**Supplementary Material**

for Manuscript “**Imaging Chromatin Interactions at near-Kilobase Resolution Via Tn5-FISH**”

**Supplementary Note S1**

**Supplementary Figures S1-S3**

**Supplementary Table S1-S2**

**Supplementary Reference**

Supplementary Note S1

**MATERIALS AND METHODS**

**Cell culture**

K562 and GM12878 cells were purchased from China Infrastructure of Cell Line Resources (Peking Union Medical College, Beijing, China) and maintained in RPMI1640 medium (Gibco, U.S.A.) supplemented with 10% fetal bovine serum (FBS, Gibco, U.S.A.), 50 units/ml penicillin and streptomycin (Gibco, U.S.A.), and Non-Essential Amino Acids (NEAA, Gibco, U.S.A.) as instructed.

**Tn5 FISH probe preparation**

Tn5-FISH probes were constructed as follows. First, sequences of interacting sites (either predicted by CMIC or previously reported) were downloaded from UCSC genome browser with repeats masked as N. DNA fragments used for probe library generation were amplified by PCR and recovered by DNA Cleanup kit (D4014, Zymo research, U.S.A.). Probe library was constructed with 50 ng DNA fragments treated by V50 Tn5 enzyme provided in Vazyme TruePrep DNA Library Prep kit (TD501, Vazyme, China) at 55℃ for 10 min, recovered and immediately amplified by Q5 enzyme (NEB, U.K.) with amplification primers as instructed by DNA Library Prep kit. The amplified probe library could serve as templates for Tn5-FISH probes. The Tn5-FISH probes were obtained by a second PCR amplification with fluorescence-tagged primers. After recovered by DNA Cleanup kit, Salmon sperm DNA (Invitrogen, U.S.A.) was added into the Tn5-FISH probes (50 μg Salmon sperm DNA per 1 μg Tn5-FISH probes), ethanol precipitated, and dissolved in DNA FISH buffer (50% deionized formamide, 10% Dextran Sulfate, 2× SSC, all from Sigma, Germany) at concentration of 20 ng Tn5-FISH probes per microliter.

For the details of all primers used, see Supplementary Table S2.

**Traditional FISH probes generated by Nick Translation of BAC clone**

Traditional FISH probes were prepared as previously described (Citation). Briefly, BAC clones were ordered from Thermo Fisher, and obtained by BACMAX kit (Epicenter, U.S.A.). For 1 μg BAC clone, 1 μL DNA Pol I and 200 U DNase I (NEB, U.K.), 1 mM each of dATP, dCTP, dGTP and 1 mM 1:2 ratio mixed fluorescence-tagged dUTP/dTTP were added together (Invitrogen, U.S.A.), vortex mixed and incubated at 15℃ for 2 h, then ethanol precipitated with 50 μg Salmon sperm DNA and 5 μg Cot1 DNA (Invitrogen, U.S.A.), and finally re-dissolved in DNA FISH buffer.

**Multi-color Tn5-FISH preparation**

The procedure of Tn5-FISH was the same as traditional FISH as previously described (Citation). Briefly, cells were fixed by 4% paraformaldehyde and permeated by 0.1% Triton-X 100, then incubated at 20% glycerol for 30 min. After 3 times quick freeze-thaw by liquid nitrogen, cells were further permeated with 0.5% Triton-X 100 and treated with 0.1M HCl. After thorough washing with PBS, cells were incubated with pre-hybridization buffer (50% deionized formamide, 2× SSC). As for two-color Tn5-FISH, each color of 10 ng Tn5-FISH probe was mixed with DNA FISH buffer and applied to cells. The FISH program was set to 75℃ for 5 min, then 37℃ overnight. On the next day, after 3 times washing with FISH-washing buffer (0.2% CA-630, 2× SSC), cells were counterstained with 2ng/mL DAPI, and sealed with Mowiol mounting medium. The slides were then imaged either by Carl Zuess LSM780 confocal microscope or Nikon A1 SIM microscope. For imaging with LSM780, samples were sequentially illuminated by 405nm, 488nm, 568nm, 594nm and 647nm lasers using 63X oil Apoplan objective lens on Axio2 microscope. For SIM imaging, samples were sequentially illuminated by 405nm, 488nm, 568nm and 647nm lasers using 100X oil Apoplan objective lens on Nikon A1 microscope.

**Image processing and Quantification of FISH signals**

The obtained Tn5-FISH images were processed by FIJI software (version 1.25h, from NIH) and the videos were processed and all images were quantified by Imaris 9.2.0 (Bitplane, Switzerland). For simple demonstration, images were deconvoluted by Huygens 17.10 (Scientific Volume Imaging b.v., Netherlands) software using default confocal model, then the Brightness/Contrast was adjusted in FIJI. Videos of 3D reconstruction were generated by Imaris 9.2.0 after adjustments of Brightness/Contrast. FISH signals were measured and quantified by Imaris 9.2.0.

**Supplementary Figures (S1-S3)**

**Figure S1**

**a**

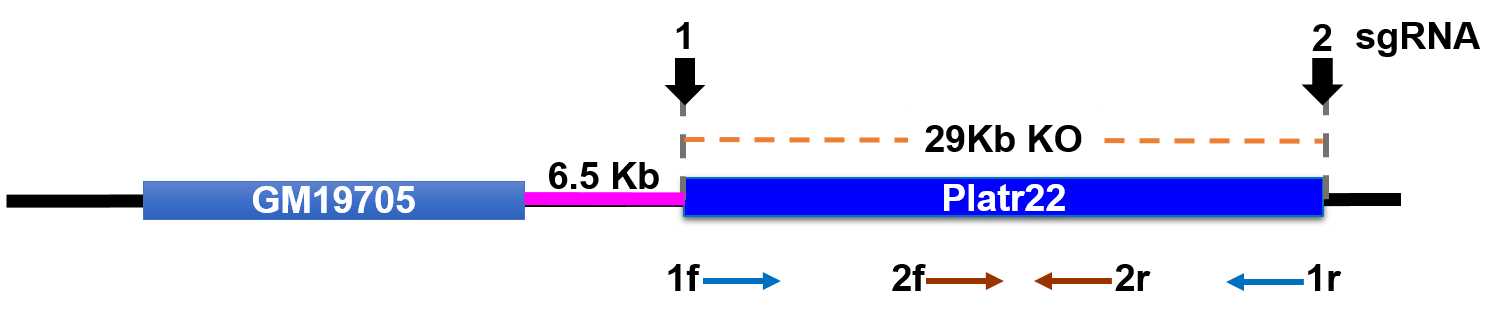

**b**

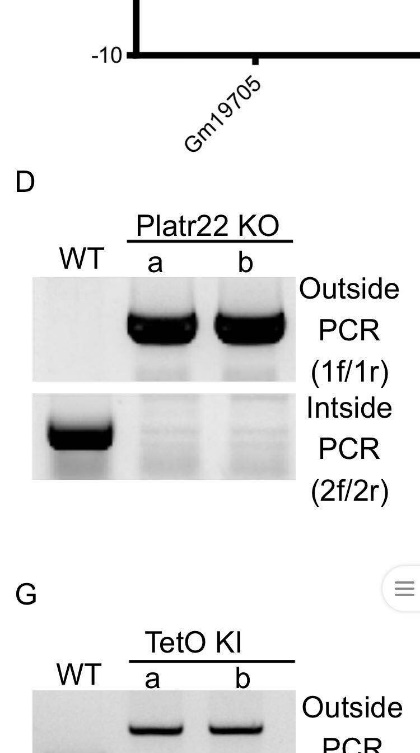

**Figure S1. CRISPR knockout of Platr22 loci (Chr1:136696211-136725728) in mESCs.**

(a) Site 1 and 2 are for sgRNA. Briefly, two sgRNAs were designed and constructed into PGL3-U6 vector. Then stable Platr22-KO mESCs were selected by puromycine after transfection. SE, super enhancer.
(b) Single clone of KO cell lines were verified by genomic PCR.

**Figure S2**

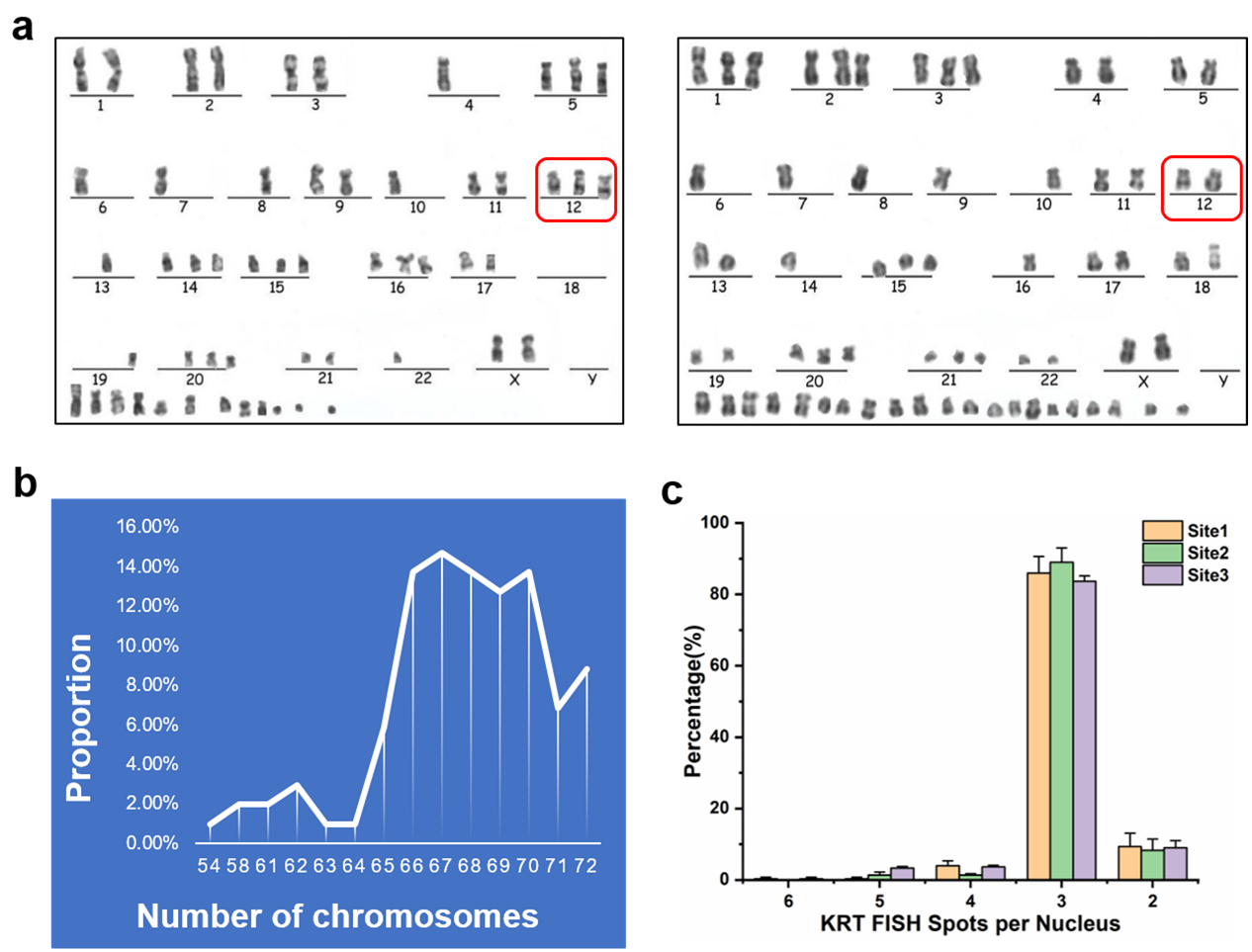

**Figure S2. The majority of K562 cells have more than two homologs per chromosome.**

(a) Karyotyping of K562 cells used in this manuscript indicated that chromosome 12 can have either two (**a**, left panel) or three (**a**, right panel) homologs.
(b) The distribution of chromosome homologues indicates that the majority of K562 cells are quasi-triploid.
(c) The number of KRT FISH spots were counted in 50 cell nuclei. More than 80% of K562 harbor three homologs, only less than 20% harbor two homologs.

**Figure S3**

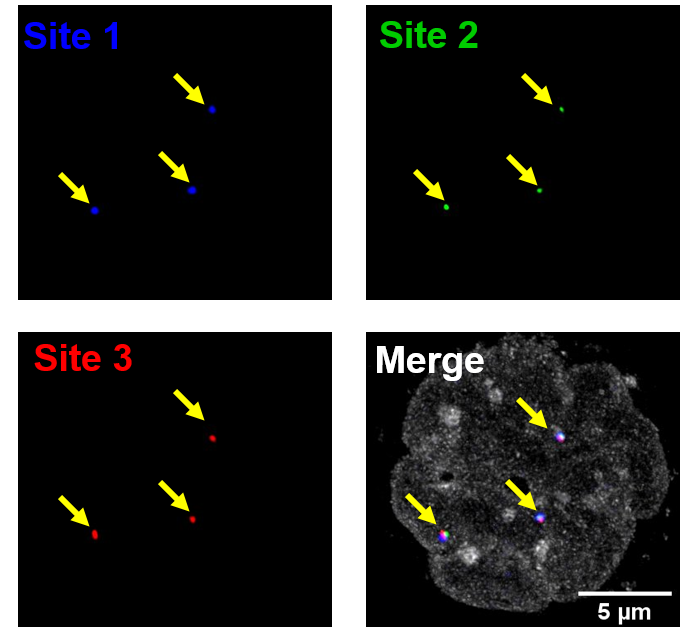

**Figure S3. The predicted chromatin interactions within KRT gene cluster II on Chr12 in K562 cells, were verified by traditional BAC FISH.** Yellow arrows indicate co-localized BAC FISH signals. The nucleus is rendered in gray for the better visualization of three-color FISH.

**Supplementary Tables (S1-S2)**

**Supplementary Table S1 Statistical summary of parameters of Tn5-FISH and traditional BAC FISH for KRT loci at chr12.**

The decrease of the probe size (from 7988 bp to 1170 bp) might lead to the decrease of the hybridization efficiency and the increase of false positive rate.

| The length of Tn5 probe target | **7988bp** | **4145bp** | **2580bp** | **1170bp** |
| --- | --- | --- | --- | --- |
| BAC FISH signals in total | 136 | 162 | 150 | 161 |
| Tn5 FISH signals in total | 144 | 172 | 138 | 169 |
| BAC-Tn5 Co-localized | 128 | 153 | 129 | 139 |
| False positive* | 16 | 19 | 9 | 30 |
| Hybridization efficiency compared to BAC** | 0.94 | 0.94 | 0.86 | 0.86 |
| False positive rate | 0.11 | 0.11 | 0.065 | 0.178 |
| Cells counted with good FISH (BAC)*** | 50 | 52 | 50 | 50 |
| Cells counted with good FISH (Tn5) | 51 | 52 | 47 | 49 |
| Total cell counted | 53 | 52 | 50 | 50 |

**Notes:**

*. False-positive is defined as Tn5-FISH spots that do not co-localize with BAC FISH spots.

**. Hybridization efficiency is defined as the ratio between the number of BAC-Tn5 spots co-localized with FISH spots and the number of total BAC FISH spots.

***. According to Fig.S6, cells with good FISH signal means there are 2-6 FISH spots (most of cells have 3 FISH signal spots) identified within the space defined by DAPI-stained nucleus in each individual cell from FOV.

Colors in the table: orange-BAC for site 1; green-BAC for site 2; blue-BAC for site 3.

**Table S2. Primers used in Tn5-FISH experiment in this manuscript**

| 1) **Primers for Fluorescence Probes** | |  |  |
| --- | --- | --- | --- |
| **Primer** | **Forward Sequence (5'to3')** | **Reverse Sequence (5'to3')** | **5' Modification** |
| TAMRA Probe | TCGTCGGCAGCGTCAGATGTGTATAAGAGACAG | GTCTCGTGGGCTCGGAGATGTGTATAAGAGACAG | TAMRA |
| 6-FAM Probe | CAGCGTCAGATGTGTATAAGAGACAG | GTCTCGTGGGCTCGGAGATGTGTATAAGAGACAG | 6-FAM |
| Cy5 Probe | TCGTCGGCAGCGTCAGATGTGTATAAGAGACAG | GGCTCGGAGATGTGTATAAGAGACAG | Cy5 |
| Amplification Primers | TCGTCGGCAGCGTCAGATGTGTATAAGAGACAG | GTCTCGTGGGCTCGGAGATGTGTATAAGAGACAG | Unmodified |
| 2) **Primers for GM19705 in mouse ESC cels** | |  |  |
| **Primer** | **Forward Sequence (5'to3')** | **Reverse Sequence (5'to3')** | **Fragment size (bp)** |
| GM19705-1 | CTTTTGCAGAGTTCAACCTGTTTGGG | CCAGAGCCAGCCTCTGTATACCC | 1593bp |
| GM19705-2 | GGGGTGTAAAACATACACAGGGGGA | CATTTTGAGAGGCCTGGATTGCTCCA | 1328bp |
| 3) **Primers for Platr22 in mouse ESC cels** | |  |  |
| **Primer** | **Forward Sequence (5'to3')** | **Reverse Sequence (5'to3')** | **Fragment size (bp)** |
| Platr22-1 | CCAGAAGACGCCAGTTCAAATTCCA | TGCAAATGTGTCATGACACACGTGT | 1967bp |
| Platr22-2 | GCAGCCTAGGCTGGTCTTGATCC | GGCCTCTCTGAGTACTGCATGTAC | 1923bp |
| 4) **Primers for KRT Sites in K562 cells** | |  |  |
| **Primer** | **Forward Sequence (5'to3')** | **Reverse Sequence (5'to3')** | **Fragment size (bp)** |
| Site1-1 | TACCATGTCAGATGAAGGGAT | TGCACAGTTTGGACAGATG | 1560 |
| Site1-2 | GTCCAGCTCACCACTAGGA | ACCGACTAAGATGCAGAATC | 2463 |
| Site1-3 | GAGAGTGATGATCCTACTTCG | GGTGTAGTTTTCAGACACCA | 1490 |
| Site1-4 | AGCTACACTTTCGAGGGTT | GCAGACATACCAATGACGGA | 2475 |
| Site2-1 | ATCCTTCCAGTGTTAGGTTGA | TTGTCAGGTCTCAACGGTCT | 2580 |
| Site3-1 | ACCAGGCTTGGCCTACTAGA | CTTCCTGGAAGAATGGTCTTC | 1170 |
| Site3-2 | CTCAGGTCTATGCCTGCATC | CATATGGTTTCTGTATGGCTCC | 1441 |
| Site3-3 | TGAGCGCCTTAGCCAGGAGT | GAAGGCACAGGGTTGGAGGT | 1534 |
| 5) **Primers for IRS1 Sites in GM12878 cells** | |  |  |
| **Primer** | **Forward Sequence (5'to3')** | **Reverse Sequence (5'to3')** | **Fragment size (bp)** |
| Site1-1 | GCTTAAACACATCCCTGGG | GGATCATCTCTTTCCCCAC | 517bp |
| Site1-2 | TAGTGGAATTGTTGCCGGAC | CCATTGCCATACAGAGAAGTTG | 689bp |
| Site1-3 | CTCTCTCTTTTTTTTTGGAAAGC | GCTCAAGGTTTCCATTTTACTC | 1895bp |
| Site1-4 | GCAAGTGGCAAAATATGGG | GGATATGGGGCATAGTTTAAGAG | 1606bp |
| Site2-1 | GTGTTATTTTTTCCACCTCTTCC | CTGTCACATTCAACCGCTC | 1682bp |
| Site2-2 | GGATGGACATAAGATAAGGAGC | CACTAGCTTTATCTCCCAGC | 596bp |
| Site2-3 | GCATGCAGCAATGTACAGAAC | CACCAGTCATTCCTATACGGC | 399bp |
| Site3-1 | GGGTATTTTTTTTAAACCCTGC | GTGCTGAGTTTTAAACAATATATTATC | 2259bp |
| Site3-2 | CCATGCTATTAACCAAAAATTTATAG | GTTGTGTTCTAAAGGGAAAATATG | 1935bp |
| Site3-3 | GAAAGAAATAGAGTCTTGTGTCC | CAGCTTTAGTGAGAAAATAAATTTG | 2013bp |

**Supplementary References:**

1.	Rowley, M.J. & Corces, V.G. *Nat Rev Genet* **19**, 789-800 (2018).

2.	Tang, Z. et al. *Cell* **163**, 1611-1627 (2015).

3.	Fang, R. et al. *Cell Res* **26**, 1345-1348 (2016).

4.	Singh, S. et al., 085241 (2018).

5.	Zeng, W. et al. *BMC Genomics* **19**, 84 (2018).

6.	He, B. et al. *Proc Natl Acad Sci U S A* **111**, E2191-2199 (2014).

7.	Roy, S. et al. *Nucleic Acids Res* **43**, 8694-8712 (2015).

8.	Whalen, S. et al. *Nat Genet* **48**, 488-496 (2016).

9.	Zhu, Y. et al. *Nat Commun* **7**, 10812 (2016).

10.	Ibn-Salem, J. & Andrade-Navarro, M.A., 257584 (2018).

11.	Kai, Y. et al. *Nat Commun* **9**, 4221 (2018).

12.	Mikolov, T. et al. in Proceedings of the 26th International Conference on Neural Information Processing Systems - Volume 2 3111-3119 (Curran Associates Inc., Lake Tahoe, Nevada; 2013).

13.	Li, G. et al. *Nucleic Acids Res* **45**, e4 (2017).

14.	Sandelin, A. et al. *Nucleic Acids Res* **32**, D91-94 (2004).

15.	Kouzarides, T. *Cell* **128**, 693-705 (2007).

16.	Sanborn, A.L. et al. *Proc Natl Acad Sci U S A* **112**, E6456-6465 (2015).

17.	Wen, Z. et al. *Cell Biol Toxicol* **34**, 471-478 (2018).

18.	Lieberman-Aiden, E. et al. *Science* **326**, 289-293 (2009).

19.	Schones, D.E. et al. *BMC bioinformatics* **8**, 19 (2007).
